# Myoelectric Prosthesis Control using Recurrent Convolutional Neural Network Regression Mitigates the Limb Position Effect

**DOI:** 10.1101/2024.02.05.578477

**Authors:** Heather E. Williams, Ahmed W. Shehata, Kodi Y. Cheng, Jacqueline S. Hebert, Patrick M. Pilarski

## Abstract

Although state-of-the-art myoelectric prostheses offer persons with upper limb amputation extensive movement capabilities, users have not been afforded a reliable means to control common movements required in daily living. Many proposed prosthesis controllers use pattern recognition, a method that learns and recognizes patterns of electromyographic (EMG) signals produced by the user’s residual limb muscles to predict and execute device movements. Such control becomes unreliable in high limb positions—a problem known as the limb position effect. Pattern recognition often uses a classification algorithm; simple to implement, but limits user-initiated control to only one device movement at a time, at a single speed. To combat position-related control deficiencies and classification controller constraints, we developed and tested two recurrent convolutional neural network (RCNN) pattern recognition-based solutions: (1) an RCNN classification controller that uses EMG plus positional inertial measurement unit (IMU) signals to offer one-speed, sequential movement control; and (2) an RCNN regression controller that uses the same data capture technique to offer simultaneous control of multiple movements and device movement velocity. We assessed both RCNN controllers by comparing them to a commonly used linear discriminant analysis classification controller (LDA-Baseline). Participants without upper limb impairment were recruited to perform multipositional tasks while wearing a simulated prosthesis. Both RCNN classification and regression controllers showed improved functional task performance over LDA-Baseline, in 11 and 38 out of 115 metrics, respectively. This work contributes an RCNN regression-based controller that provides accurate, simultaneous, and proportional movements to EMG-based technologies including prostheses, exoskeletons, and even muscle-activated video games.

## 1 Introduction

Despite the functional capabilities offered by state-of-the-art myoelectric prostheses for use by those with transradial (below elbow) amputation, users report that these modern devices are challenging to control and offer unnatural movement qualities [1, 2]. In response to user and clinical feedback, ongoing research aims to improve overall device usability, with an emphasis on control reliability [3]. Pattern recognition is a control method that has garnered much focus in upper limb prosthesis research [4]. With such control, electromyographic (EMG) signals generated in the residual limb musculature of a user are captured by device socket electrodes, and then interpreted by a controller to predict and execute the user’s intended movements. Simply stated, a prosthetic limb moves in response to muscles deliberately contracted by its user, as coordinated by a control algorithm.

Most advanced pattern recognition-based controllers tend to use a *classification* algorithm [5], which predicts one device action (or class) at a time [4]. The resulting prosthetic limb movements, consequently, appear sequential (robot-like) and are delivered at a pre-set speed—for instance, a device that offers wrist and hand capabilities cannot inherently move these components simultaneously or with varied velocity. Another control alternative uses a *regression* algorithm. This approach can predict multiple device movements at once, each of which are proportional to muscle contraction intensity. The outcome from this approach is that users can control their prosthetic wrist and hand more intuitively and in a smooth manner using varied velocities throughout reaching and grasping actions [4].

Whether a classification or regression algorithm (known as a model) is used for control, pattern recognition-based prosthesis movement predictions are contingent on earlier-captured movement data. Prior to device use, a prosthesis user must perform a series of predetermined muscle contractions using their residual limb—known as a training routine [6]. Through training, patterns observed in the captured muscle signals are associated with corresponding device actions. A training routine should not take an excessive amount of time, particularly if it must be executed to recalibrate control during prosthesis use. Indeed, user feedback surveys report that above all else (even above improved control), upper limb prosthesis users want quick-setup control solutions [7, 8].

Classification control is often employed in prosthesis research for a number of reasons: (1) it can reliably predict discrete hand and wrist classes, which reduces model complexity [3]; (2) its simpler model form can yield fewer errors versus complex ones [3, 9]; and (3) its model training routines only require execution of static muscle contractions [10], making them straightforward to develop and quick to administer. Still, predicted movement errors are known to occur with classification control, particularly during transitions between hand and wrist classes [10]. Alternatively, regression control’s main advantage is that it offers the potential for fluid prosthetic wrist/hand movements, even during transitions [3]. To achieve such fluidity, however, users must execute complex training routines that require dynamic elicitation of EMG signals with varying intensities. Choosing classification-over regression-based control evidently comes with a trade-off: movement reliability and model simplicity, over movement fluidity.

Control reliability remains a research goal, particularly as a problem known as the “limb position effect” critically impedes device reliability [11–14]. Here, surface EMG signals change when a prosthesis or other wearable device is used in untrained limb positions. It has been established that when the physical conditions for pre-training and training are dissimilar, muscle coactivation patterns can be introduced during limb movements and incorrect decoding of a user’s intent can result [11]. Muscle coactivations are introduced even among those without amputation, normally evidenced in high limb positions [9, 11]. The limb position reliability implications to prosthesis control are well acknowledged, yet remain largely unsolved [15]. To definitively solve it, a control model would have to be trained in every conceivable limb position, but in doing so would require an excessively long training time. Instead, training routines performed in selected low, midway, and high positions have been introduced to combat the limb position effect problem, with their models yielding statistically significant control improvements [12].

Researchers have also begun to add positional sensors (worn on users’ residual limbs during model training and testing) to address the problem, and have found that inertial measurement units (IMUs) can capture pertinent limb position data [12, 14, 16–18]. Our earlier work built upon this finding and successfully combined EMG and IMU data to train deep learning pattern recognition-based controllers [9]. Specifically, we developed a new type of recurrent convolutional neural network (RCNN), intended for upper limb prosthesis control. RCNNs are a network architecture for deep learning, capable of learning directly from data and handling large amounts of multimodal data [19, 20]. This makes them well-suited for training with EMG and IMU data from multiple limb positions, without requiring researchers to supply engineered features— instead, features are learned [21–23]. RCNNs have also proven to be advantageous as they can handle the intrinsic time-varying nature of muscle signals [24]. Our work capitalized on these benefits and established that position-aware RCNN controllers show promise towards mitigating the limb position effect [9]; as corroborated by our research that focused on the movements of participants without limb difference [9, 25–27].

Our body of control research led us to consider whether a more advanced *RCNN-based classification* and/or *RCNN-based regression* controller could offer improved control and functionality over earlier pattern recognition-based approaches that focused on reliability and ease-of-implementation at the expense of movement fluidity. We aimed to recommend an RCNN-based controller that uses EMG and IMU sources of movement/positional data to effectively mitigate the limb position effect. Solving this prevalent EMG-based device control problem would make inroads towards acceptance of future control solutions—by rehabilitation clinicians and users alike.

In the work presented herein, we investigate whether an RCNN classification controller (RCNN-Class) and/or an RCNN regression controller (RCNN-Reg) might offer enhanced myoelectric prosthesis control versus a conventional linear discriminant analysis (LDA) classification counterpart. The latter is commonly used to control upper limb prostheses, and as such, is often adopted in research as a baseline for comparison to other controllers [4, 9]. This work equipped non-disabled participants with a simulated device, which has been shown to be a reasonable proxy for actual myoelectric prosthesis use [28]. With the device donned, they trained and tested either RCNN-Class or RCNN-Reg, along with an LDA baseline controller (LDA-Baseline). Participants executed functional tasks across multiple limb positions during testing, with EMG and IMU data collected. Task performance metrics analysis relied on motion capture data, control characteristics analysis was based on the prosthesis’ motor data, and participants’ control experiences were gauged from their responses to surveys. Both RCNN-Class and RCNN-Reg showed improved functional task performance over LDA-Baseline. RCNN-Reg, however, offers two fundamental control advantages to users: (1) it mitigates the limb position effect, plus (2) it reliably provides smooth and simultaneous device movements. Our RCNN regression-based findings contribute to the body of myoelectric prosthesis control research by presenting a novel and advanced control approach that provides fluid movements that more closely approximate those of an intact wrist and hand. What follows are details about this work’s experimental methods, results, and promising regression-based control outcomes.

## 2 Methods

### 2.1 Overview

In this work, we compared two RCNN-based pattern recognition controllers to an LDA classification controller baseline, which applies probability theory to discover patterns in EMG data and uses engineered features to inform control [4, 29]. We considered the control performance offered by: (1) RCNN-Class versus LDA-Baseline, and (2) RCNN-Reg versus LDA-Baseline. For our investigation, three distinct controller testing sessions were undertaken, as described below.

**A) An RCNN-Class Session** (outlined in Figure 1A) required participants to don a gesture control armband equipped with both EMG and IMU sensors, plus a simulated prosthesis. While wearing this equipment, participants performed a training routine that involved *static*, isotonic forearm muscle contractions in *four* limb positions (in doing so, they trained RCNN-Class’s model). After learning how to control the device through practice, participants performed functional tasks with motion capture data recorded.

**Fig. 1.**
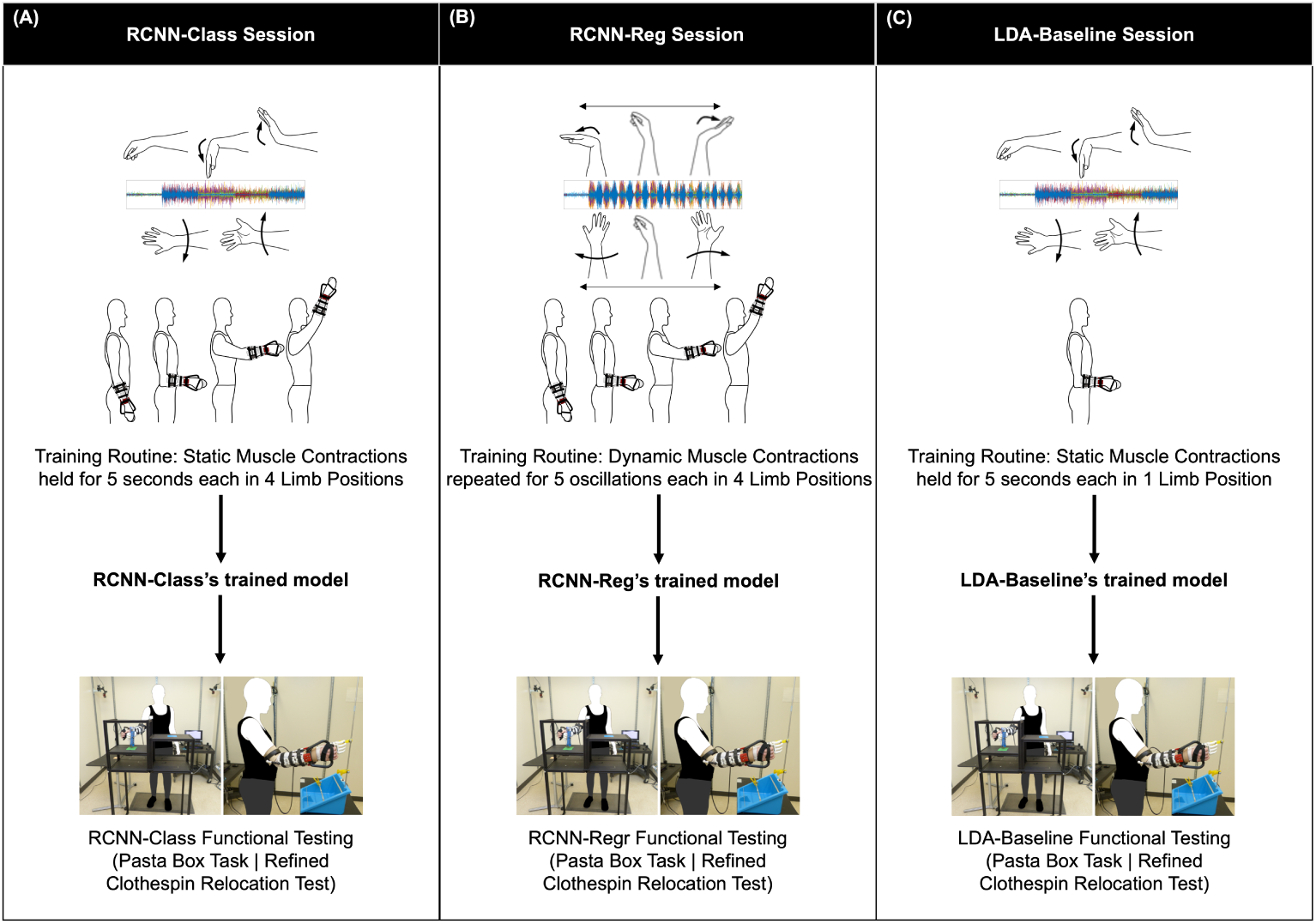
Overview of controller testing sessions: (A) RCNN-Class Session, (B) RCNN-Reg Session, and (C) LDA-Baseline Session. Note that for all training routine and testing execution, participants wore a gesture control armband (equipped with EMG and IMU sensors), along with a simulated prosthesis.

**B) An RCNN-Reg Session** (outlined in Figure 1B) required participants to don the gesture control armband and simulated prosthesis, after which they performed a training routine that involved *dynamic*, isotonic forearm muscle contractions in *four* limb positions (in doing so, they trained RCNN-Reg’s model). After learning how to control the device through practice, participants performed functional tasks with motion capture data recorded.

**C) An LDA-Baseline Session** (outlined in Figure 1C) also required participants to don the gesture control armband and simulated prosthesis, after which they performed a training routine that involved *static*, isotonic forearm muscle contractions in only *one* limb position (in doing so, they trained LDA-Baseline’s model). Notably, this training routine did not involve muscle contractions in multiple limb positions as required by the RCNN-based controllers’ models, for three reasons: (1) training in only one limb position mimics standard prosthesis controller training [6]; (2) the goal of this research was not to investigate a position-aware LDA-based controller, but rather to investigate the effectiveness of RCNN-based controllers for mitigating the limb position effect; and (3) we have previously found position-aware RCNN-based classification to better mitigate the limb position effect versus position-aware LDA-based classification [16]. After learning how to control the device through practice, participants performed the same functional tasks as used with the RCNN-based controller testing, with motion capture data recorded.

### 2.2 Participants

A total of 16 participants were recruited for this study. All participants provided written informed consent, as approved by the University of Alberta Health Research Ethics Board (Pro00086557). Each participant completed two testing sessions, with an RCNN-based controller tested in one session and LDA-Baseline tested in the other. A washout period of at least seven days was included between the two sessions to ensure that participants forgot details of their first session’s controller. The 16 participants were split into two groups.

The first group compared **RCNN-Class versus LDA-Baseline**. Eight participants were recruited for this group. They had a median age of 25 (range: 22–29) and median height of 173 cm (range: 167–181 cm), four were male, four were female, and all were right-handed. One participant had minimal previous experience with EMG pattern recognition control. Four participants in this group trained and tested RCNN-Class in their first session, whereas the other four participants trained and tested LDA-Baseline in their first session. Participants trained and tested the remaining controller in their second session, with a median of 27 days between the first and second sessions (range: 13–42 days).

The second group compared **RCNN-Reg versus LDA-Baseline**. Eight participants were recruited for this group. They had a median age of 24 (range: 19–27) and median height of 176 cm (161–193 cm), five were male, three were female, seven were right-handed and one was ambidextrous. No participants had previous experience with EMG pattern recognition control. Four participants in this group trained and tested RCNN-Reg in their first session, whereas the other four participants trained and tested LDA-Baseline in their first session. Participants then trained and tested the remaining controller in their second session, with a median of 17.5 days between the first and second sessions (range: 7–25 days).

### 2.3 Muscle Signal Data Collection

#### 2.3.1 Myo Gesture Control Armband

All participants wore a Myo gesture control armband (Thalmic Labs, Kitchener, Canada – discontinued) over their largest forearm muscle bulk [12], to facilitate capture of muscle signal data during controller training and testing. The armband was worn at approximately the upper third of participants’ right forearm, as shown in Figure 2A (with the top of the armband at a median of 28% of the way down the forearm from the medial epicondyle to the ulnar styloid process). The Myo armband contained eight surface electrodes to collect EMG data at a sampling rate of 200 Hz. It also contained one IMU to collect limb position data (three accelerometer, three gyroscope, and four quaternion data streams) at 50 Hz. Note that from the IMU, only the accelerometer data streams were used to ascertain limb position. Myo Connect software was used to stream and record EMG and IMU data in Matlab.

**Fig. 2.**
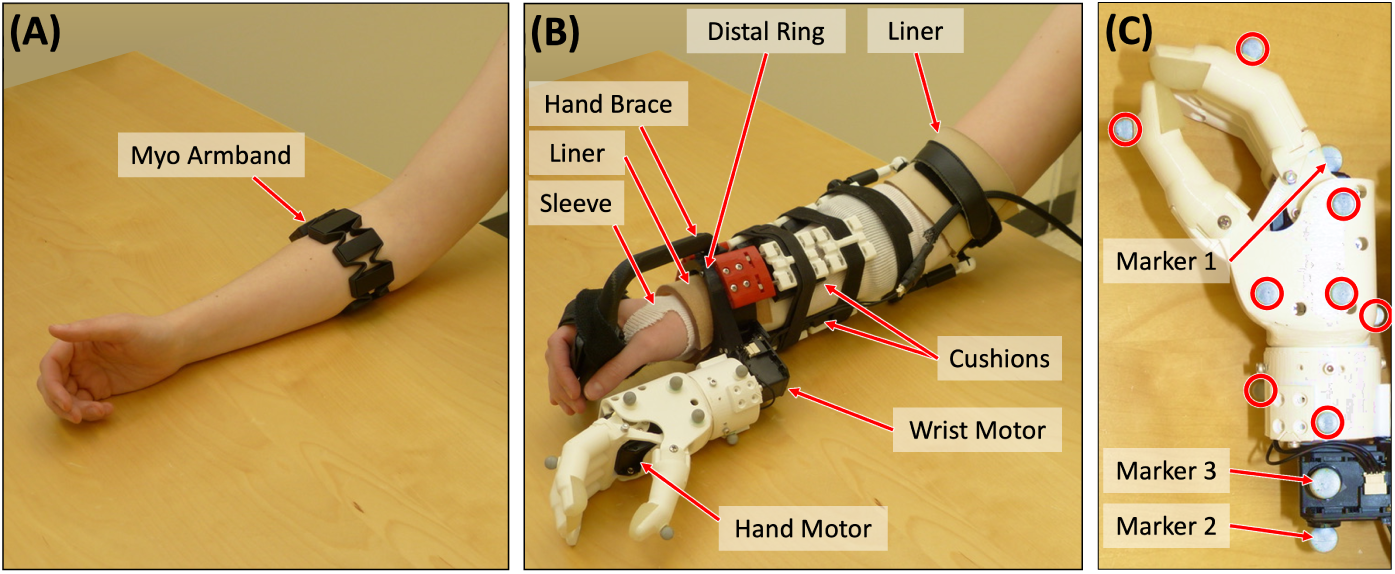
(A) Myo armband on a participant’s forearm; (B) simulated prosthesis on a participant’s forearm, with labels indicating the sleeve, two pieces of liner, hand brace, distal ring, cushions, wrist motor, and hand motor; and (C) motion capture markers affixed to the simulated prosthesis, with the eight motion capture markers that remained attached to the hand circled, and the three additional individual markers for the ski pose calibration are labelled. Adapted from Williams et al. [27].

#### 2.3.2 Simulated Prosthesis

The simulated prosthesis used in this study was the 3D-printed Modular-Adaptable Prosthetic Platform (MAPP) [30] (shown in Figure 2B). It was fitted to each participant’s right arm, to simulate transradial prosthesis use. The MAPP’s previously-published design [30] was altered to improve wearer comfort in our study—the distal ring was made to resemble the oval shape of a wrist and the hand brace was elongated so that the distal ring would sit more proximally on the wearer’s wrist. A non-proprietary 3D-printed robotic hand [31] was affixed to the MAPP beneath the participant’s hand. Wrist rotation capabilities were also added to the device. Hand and wrist movements (that is, with two degrees of freedom) were powered by two Dynamixel MX Series motors (Robotis Inc., Seoul, South Korea).

After placement of the Myo gesture control armband, participants donned a thin, protective sleeve. The MAPP was donned over the sleeve and affixed with Velcro straps. Gel-coated pieces of fabric liner (that is, with thermoplastic elastomer) were placed inside the MAPP’s distal ring and just above the participant’s elbow to ensure a comfortable fit. In addition, 3D-printed cushions, made of Ninjaflex Cheetah filament (Ninjatek, Inc.), were placed throughout the device socket (shown in Figure 2B). The MAPP was checked for secureness, a visual inspection was performed to ensure that no components were loose, and a final participant comfort check was conducted verbally.

#### 2.3.3 EMG and Accelerometer Data Processing

The EMG data from the Myo armband were filtered using a high pass filter with a cutoff frequency of 20 Hz (to remove movement artifacts), as well as a notch filter at 60 Hz (to remove electrical noise). The accelerometer data streams were upsampled to 200 Hz (using previous neighbour interpolation) to align them with the corresponding EMG data. Data were then segmented into windows (160-millisecond with a 40-millisecond offset).

### 2.4 Controller Descriptions: Model Architectures, Training Routines, & Implementation

Each of the RCNN-based controllers and their LDA-Baseline counterpart are described below. The two RCNN-based controller descriptions include details about its model’s architecture, training routine, hyperparameters used, and number of weights. LDA-Baseline’s description includes details about its statistical model implementation, training routine, features used, and number of coefficients. For each of the RCNN-based controllers presented, all hyperparameters were determined using Bayesian optimization—this includes the number of convolution layers, number of filters, filter size, pooling size, and early stopping criteria of the model. Each model’s optimization process was performed in two steps. First, an initial broad range of values were used to optimize each hyperparameter. Thereafter, values were refined using a narrower range, centered at earlier optimized values. Given that each model’s hyperparameters were optimized separately, their resulting architectures had a unique number of such with different training durations to achieve their best possible accuracies—that is, the models did not require the same number of weights nor training times to each achieve this.

#### 2.4.1 RCNN-Class Controller

##### Model Architecture

RCNN-Class’s model architecture consisted of 23 layers, as illustrated in Figure 3A. In this model, a sequence input layer first received and normalized the training data. Then, a sequence folding layer was used, allowing convolution operations to be performed independently on each window of EMG and accelerometer data. This was followed by a block of four layers: a 2D convolution, a batch normalization, a rectified linear unit (ReLU), and an average pooling layer. This block of layers was repeated twice more. Each of the three average pooling layers had a pooling size of 1x4. A block of three layers followed: a 2D convolution, a batch normalization, and a ReLU layer. The optimal number of filters in the convolution layers were determined to be 16, 32, 64, and 8, respectively, and each had a filter window size of 1x4. The next layers included a sequence unfolding layer (to restore the sequence structure), a flatten layer, a long short-term memory (LSTM) layer, and a fully connected layer. Finally, a softmax layer and classification layer yielded the final class predictions. RCNN-Class’s model had a total of 76,482 weights. Note that to prevent model overfitting, an early stopping patience hyperparameter (criteria) was set. In doing so, model training would automatically stop when the validation loss (calculated every 50 iterations) increased four times. This overfitting mitigation method chosen for this control model, is similar to that used in other works [23].

**Fig. 3.**
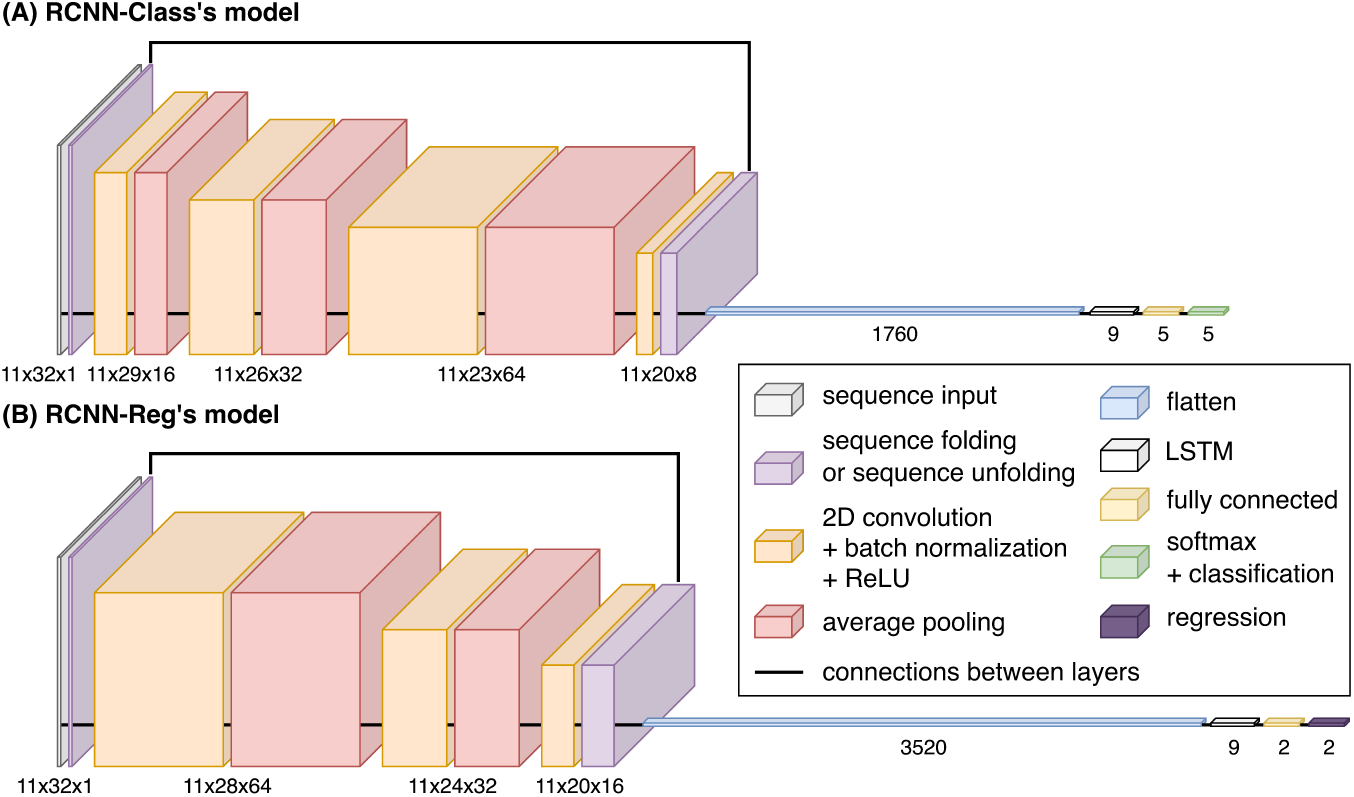
Architecture of (A) RCNN-Class’s model and (B) RCNN-Reg’s model, including the size of each layer and connections between layers.

##### Model Training Routine

Participants followed onscreen instructions, performing static muscle contractions in five wrist positions (rest, flexion, extension, pronation, and supination; shown in Figure 1A), for five seconds each. The muscle contractions were performed twice in four limb positions: arm at side, elbow bent at 90°, arm straight out in front at 90°, and arm up 45° from vertical (shown in Figure 1A). This multi-limb-position training routine was similar to those used in other real-time control studies aiming to mitigate the limb position effect [9, 12, 17]). It took approximately 200 seconds. The resulting EMG and accelerometer data, plus corresponding classes of muscle contractions, were used to train RCNN-Class’s model. A median total of 4,836 unique samples (where samples are defined as windows of 8 EMG and 3 accelerometer channels by 32 time stamps) were used for model training. Model training took a median of 5,275 iterations (meaning that a total of 25.5 million samples were observed), which required a median of 104.8 seconds to complete (with computer specifications detailed in the Control Model Implementation section).

#### 2.4.2 RCNN-Reg Controller

##### Model Architecture

RCNN-Reg’s model architecture consisted of 18 layers, as illustrated in Figure 3B. The first layers of this model are a sequence input layer and a sequence folding layer. These were followed by a block of four layers: a 2D convolution, a batch normalization, a ReLU, and an average pooling layer. This block of layers was repeated once more. Each of the two average pooling layers had a pooling size of 1x4. A block of three layers followed: a 2D convolution, a batch normalization, and a ReLU layer. The optimal number of filters in the convolution layers were determined to be 64, 32, and 16, respectively, and each had a filter window size of 1x5. The next layers included a sequence unfolding layer (to restore the sequence structure), a flatten layer, an LSTM layer, and a fully connected layer. Finally, a regression layer yielded the final predictions. RCNN-Reg’s model had a total of 140,556 weights. Note that to prevent model overfitting, an early stopping patience hyperparameter (criteria) was set. In doing so, model training would automatically stop when the validation loss (calculated every 50 iterations) increased once.

##### Model Training Routine

Participants followed onscreen instructions, performing three types of muscle contractions (shown in Figure 1B):

1. The wrist was held at rest for five seconds;
2. Dynamic muscle contractions, oscillating five times between full wrist flexion (that is, with a strong contraction that did not introduce discomfort) and corresponding full wrist extension; and
3. Dynamic muscle contractions, oscillating five times between full forearm pronation and full forearm supination.

All such muscle contractions were performed twice in four limb positions: arm at side, elbow bent at 90°, arm straight out in front at 90°, and arm up 45° from vertical (shown in Figure 1B). This multi-limb-position routine was developed in our previous offline research [9] and took approximately 300 seconds. Note that although this training routine was longer than that used for RCNN-Class, it is comparable as it allowed the model to be exposed to intermediate points between full muscle contractions. Rather than employing classes of muscle contractions in this work, two arrays with values between -1 and 1 were used to represent the flexion-extension and the pronation-supination degree of freedoms, respectively. For the static rest muscle contractions, both arrays contained zeros. For the flexion-extension oscillations, the first array contained sinusoidal values between -1 (representing full flexion) and 1 (representing full extension), and the second array contained zeros. For pronation-supination oscillations, the first array contained zeros and the second array contained sinusoidal values between -1 (representing full pronation) and 1 (representing full supination). The resulting EMG and accelerometer data, plus corresponding values, were used to train RCNN-Reg’s model. A median total of 7,420 unique samples were used for model training. Model training took a median of 500 iterations (meaning that a total of 3.7 million samples were observed), which required a median of 21.4 seconds to complete.

#### 2.4.3 LDA-Baseline Controller

##### Model Details

Four commonly used EMG features were chosen for implementation of this baseline classifier’s model: mean absolute value, waveform length, Willison amplitude, and zero crossings [32]. These features were calculated for each channel within each window of EMG data. A pseudo-linear LDA discriminant type was used, given that columns of zeros were occasionally present in some classes for some features (including Willison amplitude and zero crossings). LDA-Baseline’s model had a total of 330 coefficients.

##### Model Training Routine

Participants followed onscreen instructions, performing muscle contractions in five wrist positions (shown in Figure 1C), for five seconds each. The muscle contractions were performed twice, with the participants’ elbow bent at 90° (shown in Figure 1C). This single-position routine mimicked standard myoelectric prosthesis training [11] and took approximately 50 seconds. The resulting EMG data and corresponding classes of muscle contractions were used to train LDA-Baseline’s model. Features calculated from a median total of 1,210 unique samples (where samples are defined as windows of 8 EMG channels by 32 time stamps) were used for model training. Model training required a median of 0.7 seconds to complete.

#### 2.4.4 Control Model Implementation

Each of the three control models implemented in this study was trained using Matlab software running on a computer with an Intel Core i9-10900K processor (3.70 GHz), a NVIDIA GeForce RTX 2080 SUPER graphics card with 8GB GDDR6, and 128 GB of RAM. RCNN-Class’s, RCNN-Reg’s, and LDA-Baseline’s models were trained in median times of 104.8, 21.4, and 0.7 seconds, respectively. After completion of all model training, the simulated prosthesis was programmatically controlled as follows:

1. Matlab code was written to receive EMG and accelerometer data, from which the controller predicted intended wrist movements. Note that for RCNN-Reg, additional code was written to smooth predictions using a moving average filter (averaging the current prediction with the previous prediction) [9, 33].
2. Matlab code was also written to enable predicted motor instruction transmission to the device wrist/hand, based on resulting classifications—that is, via brachI/O-plexus software [34], where received flexion signal data were translated to hand close, extension to hand open, pronation to move wrist in a counter-clockwise rotation, and supination to move wrist in a clockwise rotation. Note that for RCNN-Reg, small prediction values were suppressed to 0 as required [9, 35], with a median value of ±0.05 (range: 0.02–0.1), out of a total signal range of -1 to 1.
3. brachI/Oplexus relayed the corresponding instructions to the simulated prosthesis’ motors. The motor instructions and positions were recorded with a sampling rate of 50 Hz.

### 2.5 Simulated Device Control Practice & Testing Eligibility

Recall that for each of the RCNN-Class, RCNN-Reg, and LDA-Baseline testing sessions, participants wore a gesture control armband (equipped with EMG and IMU sensors), along with a simulated prosthesis. With all such equipment donned and during each testing session, participants took part in 40-minute control practice periods. During this period, they were taught how to operate the simulated prosthesis using isometric muscle contractions, under three conditions:

1. Controlling the hand open/close while the wrist rotation function was disabled. They practiced grasping, transporting, and releasing objects at varying heights.
2. Controlling wrist rotation while the hand open/close function was disabled. They practiced rotating objects at varying heights.
3. Controlling the hand open/close function in concert with the wrist rotation function. They practiced tasks that involved grasping, transporting, rotating, and releasing objects at varying heights.

For RCNN-Reg control practice, each of the three conditions also required participants to practice controlling device movement velocity (that is, trying to slow down and speed up device movements). Furthermore, Condition 3 included an opportunity to practice simultaneous control of the two degrees of freedom (that is, performing wrist rotation and hand open/close at the same time).

Following all practice periods, participants were tested to determine whether they could reliably control the simulated prosthesis. For such testing, two cups were situated in front of them at two different heights, with a rubber ball in one of the cups. Participants were asked to pour the ball between the two cups, and instances when they dropped the ball or a cup were recorded. If participants could not complete at least 10 pours with a success rate of at least 75% within 10 minutes, the session was ended, and they were removed from the study. All participants passed this test.

### 2.6 Motion Capture Setup & Kinematic Calibrations

After participants were deemed eligible for controller testing, the following motion capture steps were undertaken.

#### Step 1: Motion Capture Setup

A 10-camera OptiTrack Flex 13 motion capture system (Natural Point, OR, USA) was used to capture participant movements and task objects at a sampling rate of 120 Hz. Eight individual markers were placed on the simulated prosthesis hand, circled in Figure 2C (one on the thumb, one on the index finger, and the remaining six throughout the back and side of the hand to ensure reliable rigid body tracking). Rigid marker plates were also placed on each participant’s right forearm (affixed to the simulated prosthesis socket), upper arm, and thorax, in accordance with Boser et al.’s cluster-based marker model [36].

#### Step 2: Kinematic Calibrations

Each participant was required to perform two kinematics calibrations. As per Boser et al., the first calibration called for participants to hold an anatomical pose [36], for capture of the relative positions of the hand markers and motion capture marker plates when wrist rotation and shoulder flexion/extension angles were at 0°. The second calibration required participants to hold a ski pose [36], for the purpose of refining wrist rotation angles. Here, three additional individual markers were affixed to the simulated prosthesis, as shown in Figure 2C:

1. One marker placed on the top of the prosthesis’ hand motor, with the device hand closed
2. One marker placed on the bottom of the prosthesis’ wrist motor, forming a line with the first marker (to represent the axis about which the wrist rotation occurred)
3. One marker placed on the side of the prosthesis’ wrist motor (to create a second axis, perpendicular to the axis of wrist rotation)

Upon completion of the two kinematics calibrations, all Step 2 markers were removed. What remained were only those markers affixed during Step 1 for data collection purposes.

### 2.7 Controller Testing

Testing required participants to execute functional tasks that mimicked activities of daily living across multiple limb positions, with motion capture data and participant survey results as outputs.

#### 2.7.1 Functional Task Execution

Motion capture data were collected while participants executed the following functional tasks.

##### Pasta Box Task (Pasta)

Participants were required to perform three distinct movements, where they transported a pasta box between a 1^st^, 2^nd^, and 3^rd^ location (a side table and two shelves at varying heights on a cart, including across their midline) [37]. The task setup is shown in Figure 4A. Motion capture markers were placed on all task objects, as per Valevicius et al. [37]. Participants performed a total of 10 Pasta trials. If participants dropped the pasta box, placed it incorrectly, performed an incorrect movement sequence, or hit the frame of the task cart, the trial was not analyzed. Pasta was the first of two functional tasks performed as it was considered easier.

**Fig. 4.**
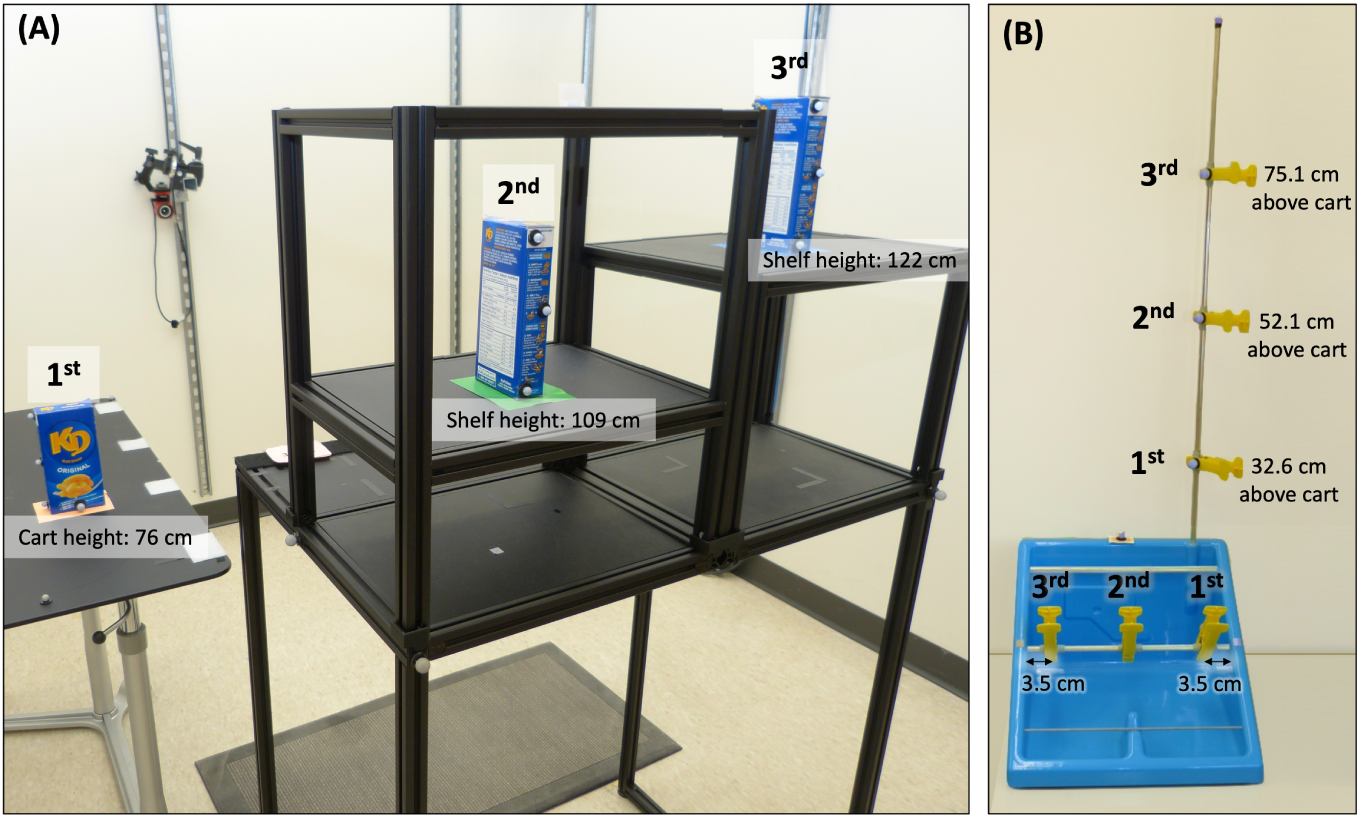
Task setup for (A) Pasta and (B) RCRT Up and Down, including task dimensions. In panel (A), the 1^st^, 2^nd^, and 3^rd^ pasta box locations are labelled. The pasta box movement sequence is 1^st^ →2^nd^ →3^rd^ →1^st^ locations. In panel (B), the 1^st^, 2^nd^, and 3^rd^ clothespin locations on the horizontal and vertical bars are labelled. The clothespin movement sequences in RCRT Up are horizontal 1^st^ →vertical 1^st^, horizontal 2^nd^ →vertical 2^nd^, and horizontal 3^rd^ →vertical 3^rd^ locations. The clothespin movement sequence in RCRT Down follows the same order, but with each clothespin moved from vertical to horizontal locations. Adapted from Williams et al. [27].

##### RCRT

Participants were required to perform three distinct movements using clothespins. They moved three clothespins between 1^st^, 2^nd^, and 3^rd^ locations on horizontal and vertical bars [38]. To simplify trial execution, RCRT was split into RCRT Up and RCRT Down trials. The task setup for these trails is shown in Figure 4B. During Up trials, participants moved the three clothespins from the horizontal bar to the vertical bar, and during Down trials, they moved the clothespins from the vertical bar to the horizontal bar. A height adjustable cart was set up—with its top surface height situated at 65 cm below the participant’s right shoulder. This setup ensured that the top of each participants’ shoulder was aligned with the midpoint between the top two targets on the vertical bar used in the RCRT Up and Down trials. Motion capture markers were placed on all task objects, as per our earlier research [26]. Participants performed a total of 10 Up trials and 10 Down trials. If participants dropped a clothespin, placed it incorrectly, or performed an incorrect movement sequence, the trial was not analyzed. Performance of RCRT Up and Down trials were alternated, beginning with RCRT Up.

#### 2.7.2 Perceived Control via Participant Survey Administration

At the end of every controller testing session, participants completed two surveys: the NASA-TLX [39] and a usability survey [40]. The former was administered using the official NASA-TLX tablet application, where participants scored their device control workload demand on a continuous rating scale with endpoint anchors of low and high. The usability survey was administered on paper, where participants marked their usability scores on a continuous rating scale with endpoint anchors of 0 and 5. In their second session, participants were not reminded of their survey responses from their first session.

### 2.8 Analysis of Controller Testing Data

Statistical data analysis was undertaken after the following steps were completed: processing of the motion capture data (with some measures estimated); finalization of estimated measures; task segmentation; and calculation of control assessment metrics.

#### 2.8.1 Motion Capture Data Processing

Motion capture data were cleaned and filtered. As per Valevicius et al. [37], grip aperture was estimated using the distance between the motion capture markers on the simulated prosthesis’ index finger and thumb. In addition, a 3D object representing the simulated prosthesis’ hand was generated using the remaining 6 hand markers, for the purpose of calculating upcoming metrics [37]. Next, using calculations modified from Boser et al. [36], wrist rotation angles were estimated using the forearm and hand markers, whereas shoulder flexion/extension angles were calculated using the upper arm and thorax motion markers.

#### 2.8.2 Finalization of Grip Aperture and Wrist Rotation Measures

The grip aperture and wrist rotation measures were finalized using data from the simulated prosthesis’ two motors. As small (yet informative) adjustments in the positions of these motors may not have been detected by the motion capture cameras, we chose not to ignore them. First, the positions of the motors were upsampled to 120 Hz using linear interpolation. Then, the estimated measures were finalized—*Grip aperture:* the motion-capture-estimated grip apertures were fitted to a trinomial curve, to facilitate transforming the hand motor data to final grip aperture measures. *Wrist rotation:* the motion-capture-estimated wrist rotation angles were fitted to a binomial curve, to facilitate transforming the wrist motor data to final wrist rotation angle measures.

#### 2.8.3 Task Segmentation

The task data were segmented in accordance with Valevicius et al. [37]. For each *task*, the data from each *trial* were first divided into distinct *movements 1, 2, and 3* based on hand velocity and the velocity of the pasta box/clothespins during transport.

##### Pasta

Movements 1, 2, and 3 differentiated between: (1) reaching for the pasta box at its 1^st^ location, grasping it, transporting it to its 2^nd^ location, releasing it, and moving their hand back to a home position; (2) reaching for the pasta box at the 2^nd^ location, grasping it, transporting it to its 3^rd^ location, releasing it, and moving their hand back to the home position; and (3) reaching for the pasta box at the 3^rd^ location, grasping it, transporting it back to the 1^st^ location, releasing it, and moving their hand back to the home position.

##### RCRT Up

Movements 1, 2, and 3 differentiated between: (1) reaching for the 1^st^clothespin at its 1^st^ horizontal location, grasping it, transporting it to its 1^st^ vertical location, releasing it, and moving their hand back to a home position; (2) reaching for the 2^nd^ clothespin at its 2^nd^ horizontal location, grasping it, transporting it to its 2^nd^ vertical location, releasing it, and moving their hand back to the home position; and (3) reaching for the 3^rd^ clothespin at its 3^rd^ horizontal location, grasping it, transporting it to its 3^rd^ vertical location, releasing it, and moving their hand back to the home position.

##### RCRT Down

Movements 1, 2, and 3 differentiated between: (1) reaching for the 1^st^ clothespin at its 1^st^ vertical location, grasping it, transporting it to its 1^st^ horizontal location, releasing it, and moving their hand back to a home position; (2) reaching for the 2^nd^ clothespin at its 2^nd^ vertical location, grasping it, transporting it to its 2^nd^ horizontal location, releasing it, and moving their hand back to the home position; and (3) reaching for the 3^rd^ clothespin at its 3^rd^ vertical location, grasping it, transporting it to its 3^rd^ horizontal location, releasing it, and moving their hand back to the home position.

Then, the data from each of the three movements were further segmented into *five phases* of Reach, Grasp, Transport, Release, and Home (note that the Home phase was not used for data analysis). Finally, *two movement segments* of Reach-Grasp and Transport-Release were defined for select metrics analysis. Six final levels for data analysis resulted: *controller* (either RCNN-Class, RCNN-Reg, or LDA-Baseline), *task* (either Pasta, RCRT Up, or RCRT Down), *trial* (1–10)*, movement* (1–3), *movement segment* (Reach-Grasp or Transport-Release), and *phase* (Reach, Grasp, Transport, or Release).

#### 2.8.4 Control Assessment Metrics Calculations

The *Suite of Myoelectric Control Evaluation Metrics*, introduced in our previous work [27], were used to compare prosthesis control resulting from RCNN-Class versus LDA-Baseline and RCNN-Reg versus LDA-Baseline. A summary of these metrics can be found in Table 1.

**Table 1.**
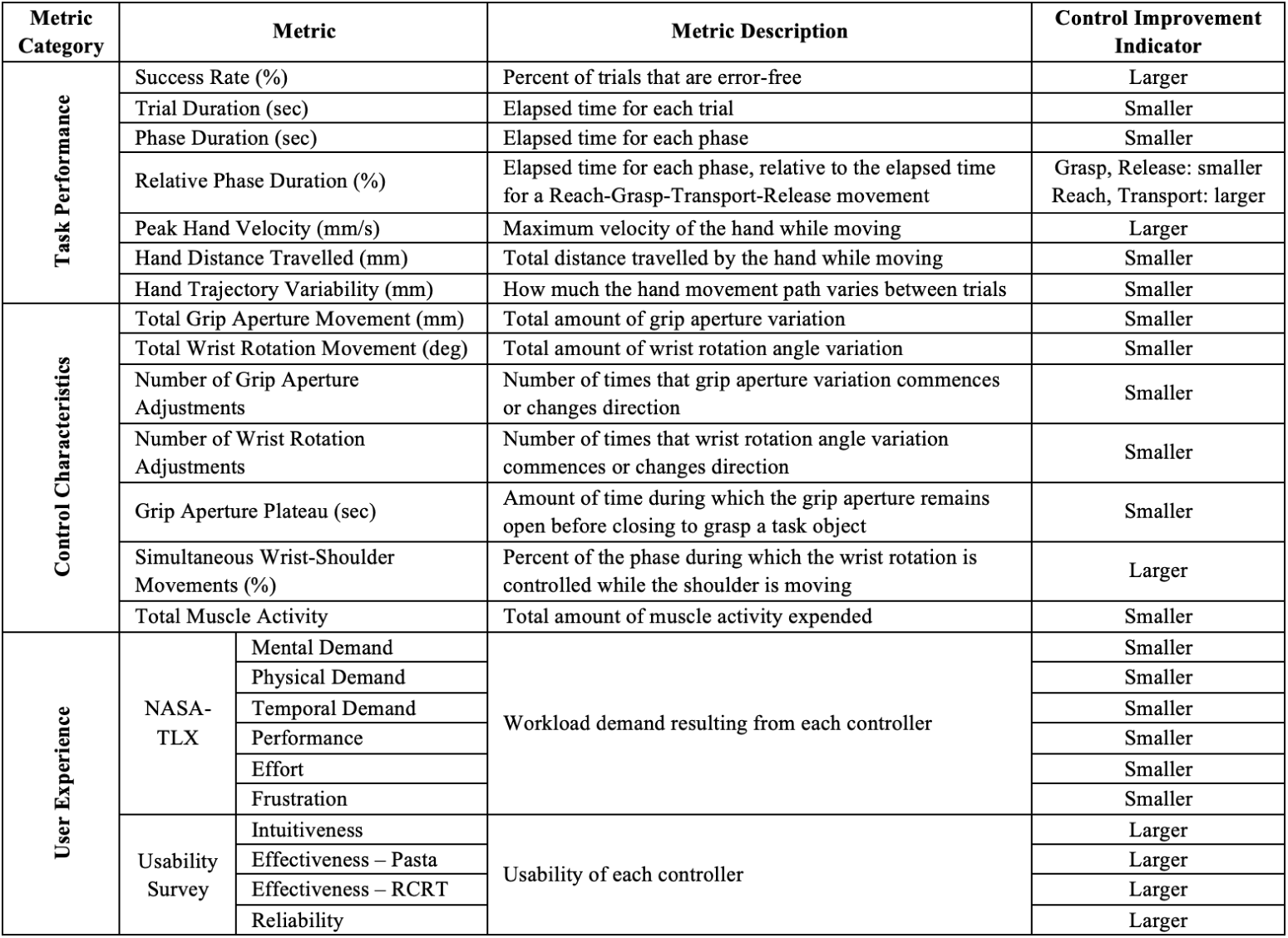
Summary of metrics used in this study, including details of whether a smaller or larger value indicates a control improvement.

To identify occurrences where the limb position effect impeded control, we analyzed our resulting Control Characteristics metrics from the suite, as outlined in our previous work [27]—the thresholds used for identification of the limb position effect remained the same, with the total muscle activity metric eliminated from analysis thereof. Specifically, this present work’s limb position effect analysis considered trends across the Control Characteristics metrics’ medians and interquartile ranges (IQRs) for each task’s three movements, with larger medians and/or larger interquartile ranges providing evidence of degraded control. Figure 5A illustrates an example where the limb position effect was identified by the increasing medians and IQRs from movement 1 (in the lowest limb position) to movement 3 (in the highest limb position). Conversely, Figure 5B illustrates an example where the limb position effect was *not* identified.

**Fig. 5.**
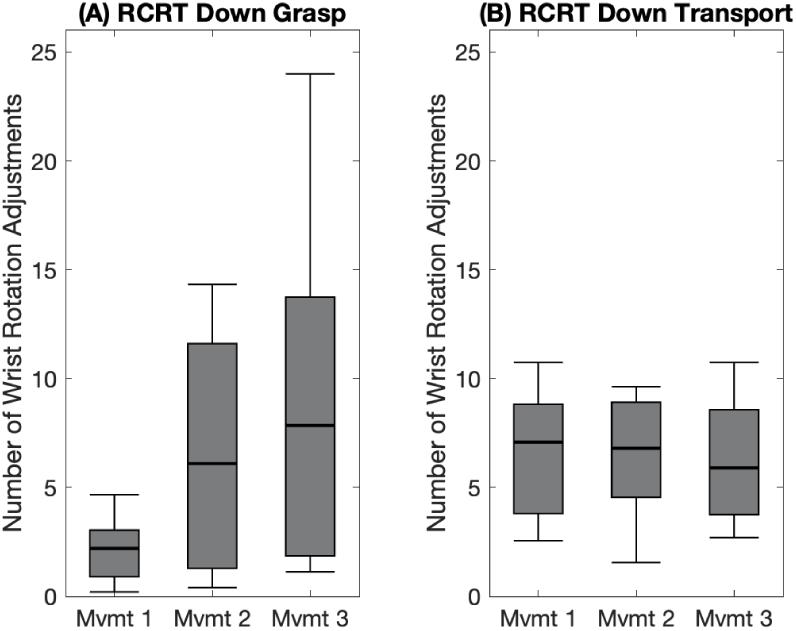
Box plots indicating LDA-Baseline number of wrist rotation adjustments in (A) RCRT Down Grasp and (B) RCRT Down Transport of each task movement (Mvmt). Medians are indicated with thick lines, and interquartile ranges are indicated with boxes.

Finally, to assess whether participants took advantage of RCNN-Reg’s simultaneous and proportional velocity control capabilities, we calculated two additional metrics from recorded motor instructions. To assess whether participants capitalized on RCNN-Reg’s **simultaneous control** capabilities, simultaneous wrist-grip movements were calculated—the percent of each phase in which the wrist rotation and the grip aperture velocities were simultaneously moving. Of note, this metric was only calculated for RCRT Up and RCRT Down tasks (given that wrist rotation movements were not necessary for Pasta). Furthermore, the metric was only applicable to Reach, Grasp, and Release phases (given that grip aperture adjustments were not necessary in Transports). To assess whether participants capitalized on RCNN-Reg’s **proportional velocity control** capabilities, instances where velocity fluctuated between 0 and full speed were counted—as indicated by the percentage of each task in which wrist rotation and grip aperture respectively varied. These two motor-recorded metrics were only calculated for RCNN-Reg, given that neither simultaneous control nor proportional velocity control were possible under RCNN-Class or LDA-Baseline alternatives.

#### 2.8.5 Statistical Analysis

To compare control resulting from RCNN-Class versus LDA-Baseline and RCNN-Reg versus LDA-Baseline, the following statistical analyses were performed using all Task Performance, Control Characteristics, and User Experience metrics:

##### For metrics that were analyzed at the phase or movement segment level

Participants’ results were first averaged across trials and movements. If results were normally-distributed, a two-factor repeated-measures analysis of variance (RMANOVA) was conducted using the factors of controller and phase/movement segment. When the resulting controller effects or controller-phase/movement segment interactions were deemed significant (that is, when the Greenhouse-Geisser corrected p value was less than 0.05), pairwise comparisons between the controllers were conducted. If results were not normally-distributed, the Friedman test was conducted. When the resulting p value was less than 0.05, pairwise comparisons between the controllers were conducted. Pairwise comparisons (t-test/Wilcoxon sign rank test) were deemed significant if the p value was less than 0.05.

##### For metrics that were analyzed at the trial level

Participants’ results were first averaged across trials, then pairwise comparisons were conducted as detailed above.

##### For metrics that were analyzed at the task or controller level

Pairwise comparisons were conducted as detailed above.

One participant’s data were not included in this study’s statistical analysis. That participant experienced difficulties with LDA-Baseline control in their second testing session (comparing RCNN-Reg versus LDA-Baseline). Although they passed their practice test, they only completed three error-free trials of Pasta and were unable to complete any error-free trials of RCRT Up or Down. Consequently, their data stemming from all controller testing sessions were excluded from analysis.

## 3 Results

The Suite of Myoelectric Control Evaluation Metrics [27] was used in this comparative controller study. The suite includes three broad categories of metrics: task performance, control characteristics, and user experience. Through an analysis of these metrics, numerous control findings were uncovered. These findings included identification of significant improvements in RCNN-based controller performance over LDA-Baseline, metrics-substantiated evidence of the limb position effect, indications that prosthesis control improvements in high limb positions can be offered by RCNN-based controllers, and that RCNN-Reg’s simultaneous and proportional velocity control capabilities yield overall improved device performance.

### 3.1 Significant Differences between RCNN-based Controllers and LDA-Baseline

Instances of significant improvements for RCNN-based control over LDA-Baseline are summarized in Figure 6—48 Task Performance and 67 Control Characteristics metrics for Pasta, RCRT Up, and RCRT Down tasks are presented. Here, 11 out of 115 metrics showed significant controller performance improvement by RCNN-Class over LDA-Baseline, and 38 out of 115 metrics showed significant controller performance improvements by RCNN-Reg versus LDA-Baseline. No significant differences between controllers were identified in User Experience metrics. All controller comparison results can be found in Appendix A.

**Fig. 6.**
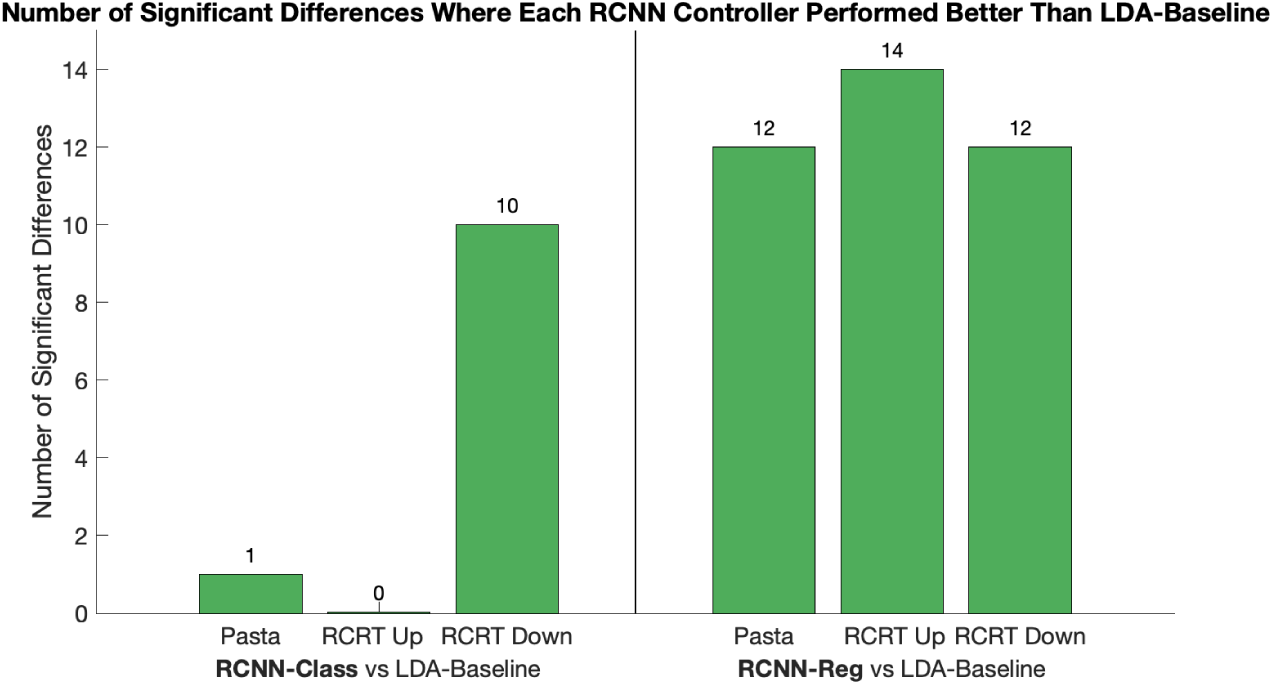
Number of significant differences between RCNN-Class versus LDA-Baseline and RCNN-Reg versus LDA-Baseline in Task Performance and Control Characteristics metrics. An average tally of performance improvement results is presented per task.

Two examples are presented below to illustrate controller outcomes during movements in high limb positions. One example presents a discrete metric, the other presents a continuous metric. Note that the LDA-Baseline results presented in these examples vary slightly between RCNN-Class versus LDA-Baseline and RCNN-Reg versus LDA-Baseline, because they are specific to two different groups of participants.

### 3.2 Discrete Metric Example: Number of Grip Aperture Adjustments

To understand whether participants altered their device control behaviour when using the simulated prosthesis in high limb positions, the Number of Grip Aperture Adjustments made by a participant in RCRT Down Grasp phases were analyzed (an example of which is presented in Figure 7). Figure 7A illustrates the median number of such adjustments for RCNN-Class versus LDA-Baseline and for RCNN-Reg versus LDA-Baseline, with significant differences identified only between the latter. To further understand differences between RCNN-Reg versus LDA-Baseline, grip aperture examples from the same participant using RCNN-Reg and LDA-Baseline control were analyzed (with Movement 3, which was at a high limb position and therefore where control was most difficult, presented in Figure 7B and 7C, respectively). In subplots 7B and 7C, the Grasp phases are highlighted, grip aperture adjustments in these phases are identified, and these adjustments are tallied. Notably, only one grip aperture adjustment was identified in the RCNN-Reg example (Figure 7B) whereas five grip aperture adjustments were identified in the LDA-Baseline example (Figure 7C). Evidently fewer adjustments were required under RCNN-Reg control.

**Fig. 7.**
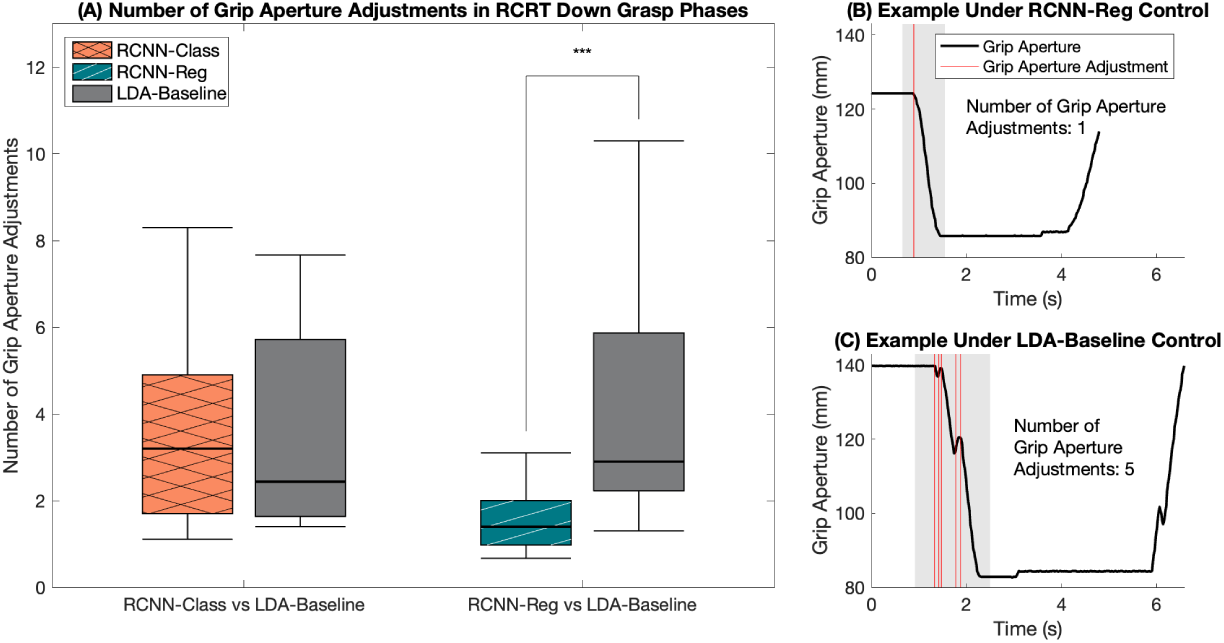
(A) Median Number of Grip Aperture Adjustments in RCRT Down Grasp phases, comparing RCNN-Class versus LDA-Baseline and RCNN-Reg versus LDA-Baseline, with interquartile ranges indicated with boxes and significant differences indicated with asterisks (*** indicates p *<* 0.05); (B) grip aperture in Movement 3 of one trial under RCNN-Reg Control from a participant; and (C) grip aperture in Movement 3 of the one trial under LDA-Baseline Control from the same participant as panel B. The Grasp phases in panels B and C are highlighted in grey and each Grasp phase grip aperture adjustment is identified with a red line.

### 3.3 Continuous Metric Example: Grip Aperture Plateaus

To further understand whether participants altered their device control behaviour when using the simulated prosthesis in high limb positions, Grip Aperture Plateaus of a participant in RCRT Down Reach-Grasp movement segments were analyzed (an example of which is presented in Figure 8). Figure 8A illustrates the median plateaus for RCNN-Class versus LDA-Baseline and for RCNN-Reg versus LDA-Baseline, with significant differences identified only between the latter. To further understand differences between RCNN-Reg versus LDA-Baseline, grip aperture examples from the same participant using RCNN-Reg and LDA-Baseline control were analyzed (with Movement 3, which was at a high limb position and therefore where control was most difficult, presented in Figure 8B and 8C, respectively). In subplots 8B and 8C, Grip Aperture Plateaus are highlighted, and these plateaus are summed. Notably, only a 0.86 s plateau was identified in the RCNN-Reg example (Figure 8B) whereas a 2.61 s plateau was identified in the LDA-Baseline example (Figure 8C). Evidently more natural movements resulted under RCNN-Reg control.

**Fig. 8.**
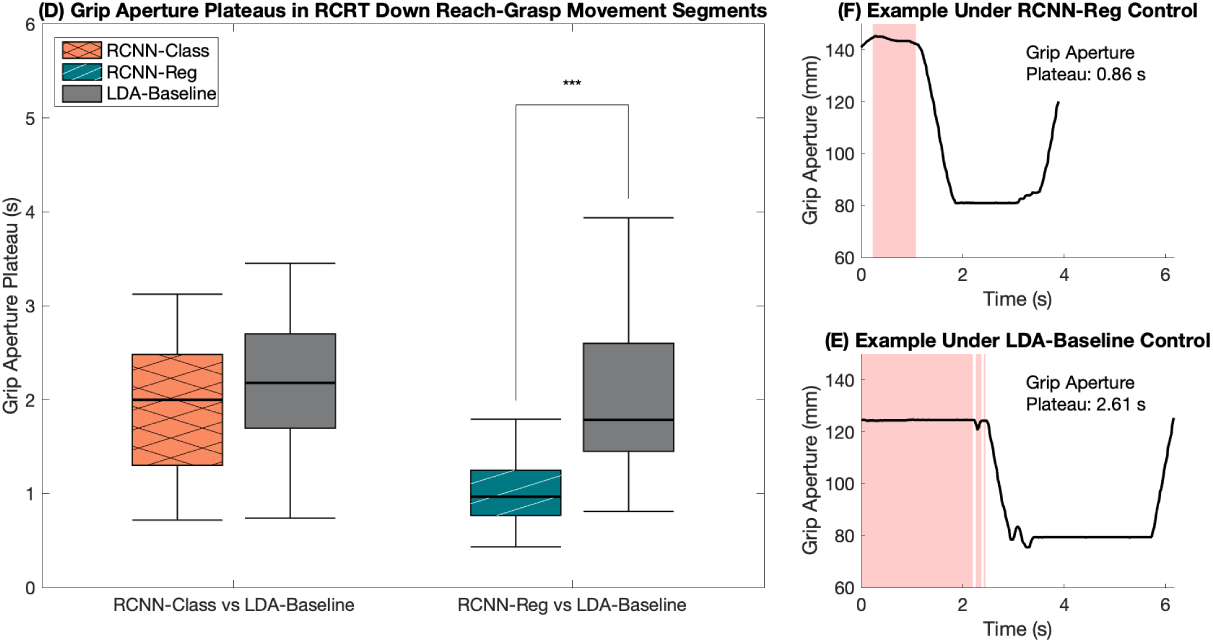
(A) Median Grip Aperture Plateaus in RCRT Down Reach-Grasp movement segments, comparing RCNN-Class versus LDA-Baseline and RCNN-Reg versus LDA-Baseline, with interquartile ranges indicated with boxes and significant differences indicated with asterisks (*** indicates p *<* 0.05); (B) grip aperture in Movement 3 of one trial under RCNN-Reg Control from a participant; and (C) grip aperture in Movement 3 of the one trial under LDA-Baseline Control from the same participant as panel B. The Grip Aperture Plateaus are identified with red shading.

### 3.4 Metrics-Substantiated Evidence of Limb Position Effect

#### LDA-Baseline

Under LDA-Baseline control, a total of 13 instances of the limb position effect were identified in Pasta and RCRT Down tasks’ Control Characteristics metrics. During Pasta execution, the following instances confirmed the limb position effect problem: in 1 of 4 metrics in Reach, 1 of 1 metric in Reach-Grasp, 4 of 4 metrics in Grasp, 2 of 4 metrics in Transport, and 2 of 3 metrics in Release. During RCRT Down execution, 3 of 4 metrics in Grasp showed evidence of the limb position effect problem (one of which is illustrated in Figure 5A). Notably, the limb position effect was not identified in RCRT Up under LDA-Baseline control.

#### RCNN-based Controllers

The limb position effect was never identified under RCNN-Class or RCNN-Reg control, suggesting that both such controllers might have mitigated the limb position effect.

### 3.5 RCNN-Reg Simultaneous and Proportional Velocity Control Capabilities

Participants in this study took advantage of the simultaneous and proportional velocity control capabilities offered by RCNN-Reg. Figure 9A illustrates simultaneous control use in Reach, Grasp, and Release phases of RCNN-Reg in RCRT Up and Down tasks. Figure 9B illustrates proportional velocity control use of grip aperture and wrist rotation using RCNN-Reg in all three tasks.

**Fig. 9.**
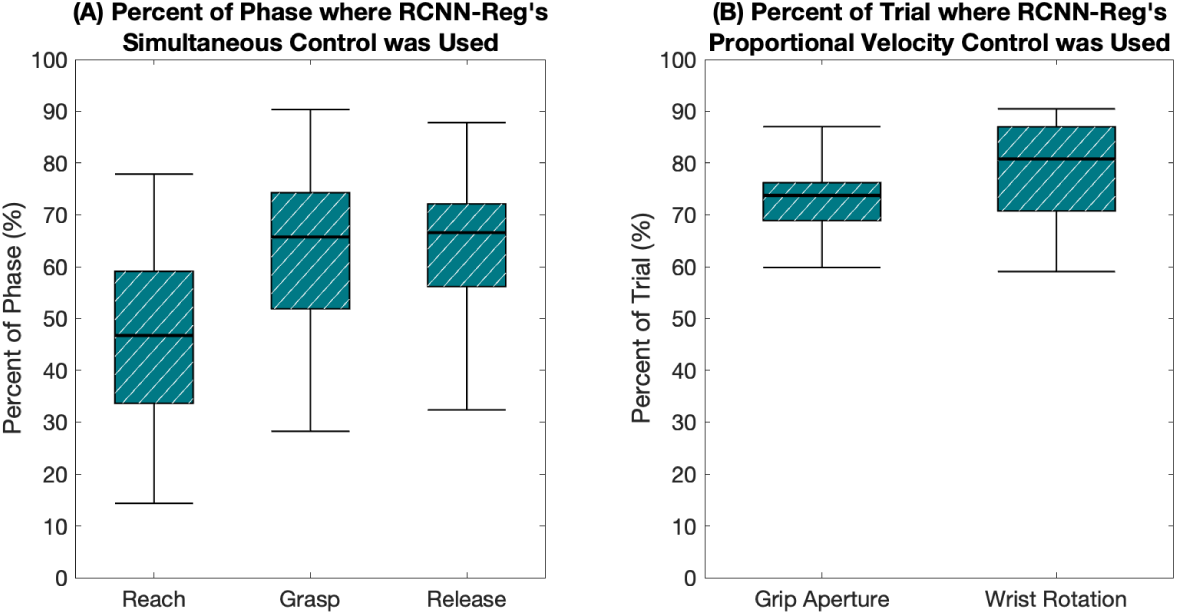
(A) The median percentage of each Reach, Grasp, and Release phase of RCRT Up and Down where grip aperture and wrist rotation were simultaneously controlled using RCNN-Reg, and (B) the median percentage of each trial where grip aperture and wrist rotation were each controlled at a velocity between no movement and maximum velocity.

## 4 Discussion

In this work, RCNN control using either classification (RCNN-Class) or regression (RCNN-Reg), versus LDA-Baseline classification control were compared. Analysis of comprehensive myoelectric control evaluation metrics [27] yielded rich, limb-position-related outcomes. What follows is a discussion about such outcomes, with a focus on understanding control challenges experienced during participants’ execution of the Pasta Box Task (Pasta) and the Refined Clothespin Relocation Test (RCRT Up and RCRT Down).

The first major finding from this comparative controller research study is that RCNN-Reg offers more accurate and position-aware prosthesis control versus LDA-Baseline. This assertion is evidenced by four key findings:

- RCNN-Reg performed significantly better than LDA-Baseline in 38 control evaluation metrics.
- Fifteen of these 38 metrics were in the specific phases where instances of the limb position effect occurred (that is, in all phases of the Pasta functional task, and Grasp phases of RCRT Down task). As such, RCNN-Reg successfully mitigated these instances of the limb position effect.
- Furthermore, RCNN-Reg performed significantly better than LDA-Baseline in every phase of each functional task performed in this study. Given this, RCNN-Reg can be said to provide accurate control under a variety of limb positions.
- Finally, RCNN-Reg performed significantly better than LDA-Baseline in 12 Pasta task metrics. Recall that this task requires limb positions not included in RCNN-Reg model’s training routine (that is, cross-body and away-from-body movements). This suggests that RCNN-Reg can maintain its predictive movement accuracy in limb positions for which it is not specifically trained.

A second major takeaway from this study is that RCNN-Reg likely offers more accurate control as compared to RCNN-Class. Although RCNN-Reg and RCNN-Class were not directly compared in this work to respect participants’ time, their respective number of significant improvements over LDA-Baseline can be compared to offer evidence of this conclusion:

- In the Pasta task, RCNN-Class was significantly better than LDA-Baseline in 1 metric and RCNN-Reg was significantly better than LDA-Baseline in 12 metrics →RCNN-Reg yielded 11 more control improvement instances.
- In RCRT Up, RCNN-Class was never significantly better than LDA-Baseline, whereas RCNN-Reg was significantly better than LDA-Baseline in 14 metrics →RCNN-Reg yielded 14 more control improvement instances.
- In RCRT Down, RCNN-Class was significantly better than LDA-Baseline in 10 metrics and RCNN-Reg was significantly better than LDA-Baseline in 12 metrics →RCNN-Reg yielded 2 more control improvement instances.

A third major finding from this work is that RCNN-Reg offers more natural functionality versus LDA-Baseline or even RCNN-Class. RCNN-Reg affords users the ability to control multiple device movements at a given time (such as those of the wrist and hand), and to control device movement velocity. It is recognized in the literature that regression-based myoelectric control is natural and flexible—device functions can be accessed in combination, with their velocities being independently controlled [41]. This work examined whether participants took advantage of these control capabilities, as offered by RCNN-Reg. The following two observations were uncovered:

- Participants *did* take advantage of simultaneous control. Under RCNN-Reg control, simultaneous movements during task execution were indeed evidenced (see Figure 9A).
- Participants also took advantage of movement velocity control capabilities. Under RCNN-Reg control, mid-velocity hand and wrist movements were evidenced during task execution (that is, velocities ranging between no movement and maximum velocity were found; see Figure 9B).

Despite the improved control outcome takeaways, particularly as offered by RCNN-Reg, no significant differences between controllers under investigation were identified in the NASA-TLX and usability surveys. Interestingly, though not statistically significant, participants did rate the mental demand required of RCNN-Reg to be high (see Appendix A). Mental demand had the largest difference between RCNN-Reg and LDA-Baseline scores, in comparison to all survey dimensions. This outcome aligns with the notion that progressive learning needs to be allocated for complicated hand-tasks, given that such tasks are mentally demanding [41]. As RCNN-Reg control offers both simultaneous and velocity control capabilities, it is reasonable to assume that introducing a revised and perhaps longer, progressive learning approach during controller practice sessions will improve users’ appreciation of these improvements. Our earlier research, which employed the same surveys, offered other important considerations—that users’ perceptions of poor or diminished control between sessions might change over time, and that reminders of earlier session control perceptions would have helped to establish first-session anchor scores [42]. This work concurs with these considerations and recommends that the Prosthesis Task Load Index (PROS-TLX) [43] be considered in future comparative prosthesis control research.

It is understood in the literature that two key factors influence user-assessments of upper limb prosthetic devices (hands in particular)—intuitive control experiences and minimal burden placed on the user (including training burden) [7, 8]. This study’s RCNN-Reg offered control improvements over RCNN-Class and LDA-Baseline but required longer training routines (training routine durations of 300, 200, and 50 seconds, respectively). It is worth noting that improved control should not come at the cost of a burdensome training routine, as the latter will overshadow users’ perceptions of control benefits [10]. The longer training routine required of RCNN-Reg, therefore, must not be ignored despite the users’ benefits of simultaneous control over multiple degrees of freedom and movement velocity control. It is recommended that routines based on continuous hand movements be streamlined, to make model training less of a burden for users. In addition, transfer learning methods should be explored as a means of minimizing training burden introduced by RCNN-Reg [26].

This current study’s RCNN-Reg success is believed to be largely due to how the training routine was implemented (despite its length) and how the prediction smoothing was undertaken. Participants followed on-screen sinusoids during training, to ensure that they executed well-paced continuous wrist and hand motions. This on-screen method kept them engaged and focused throughout all multi-limb-position training. Routine implementation methods are important, given that successfully training a control model in multiple limb positions is known to improve prosthesis control [12, 44, 45]. Along with use of an engaging multi-limb-position training method, prediction smoothing was rigorously undertaken in this study. It is recognized that smoothing a continuous model’s output, including that from RCNN-Reg, can improve model stability [46]. So together, the model training and prediction smoothing methods used in this work were key factors in RCNN-Reg’s performance success, its ability to mitigate the limb position effect, and its capacity to offer simultaneous wrist/hand movements with variable velocity.

Future work should build upon the regression-based control successes uncovered in this study, including: introducing simultaneous wrist and hand movements to the model training routine; extending model training and testing to offer more degrees of freedom; averaging three or more predictions (rather than two, as used in this work) to potentially improve prediction smoothing; and investigating the window size and overlap used in data processing (given that RCNN-based models are not yet the norm in prosthesis control). Finally, to corroborate our finding that the limb position effect can indeed be mitigated, future control model training and testing should be conducted by persons with amputation.

## 5 Conclusion

This work contributes to upper limb myoelectric prosthesis research by offering a novel regression-based controller that employs deep learning methods. The controller provides both simultaneous control of multiple device movements at a given time and control over device movement velocity, across numerous limb positions—capabilities much closer to the fluid movements of an intact wrist and hand. In this study, an RCNN classification controller and an RCNN regression controller were each compared to a commonly used LDA classification (baseline) alternative. Of these, the regression-based approach offered the most reliable and fluid prosthetic wrist/hand movements. Offering such position-aware control, however, required a training routine that was longer than that of baseline pattern recognition, and therefore could be considered burdensome to the user. Nevertheless, regression-based solutions should continue to be investigated in future RCNN control studies, as we expect newer models to yield similarly smooth and natural wrist and hand movements. We recommend that model architectures be reimagined, training routines be altered to include simultaneous wrist and hand movements, and that transfer learning methods be applied to minimize training burden. Given that RCNN regression-based control considers the complex and dynamic process of a wrist and hand functioning in concert with varied velocities, we recognize this as a promising direction for myoelectric upper limb device control that satisfies user needs and wants. As such, next-step testing and validation of RCNN regression-based control should include persons with transradial amputation, to confirm the validity of our proposed controller’s benefits. Finally, as EMG-based controllers are not restricted to upper limb prosthesis applications, this research has far-reaching implications—towards use in rehabilitation exoskeletons and even EMG-activated video games. When the limb position effect is solved, acceptance of future device movement technologies by clinicians and users should become a reality.

## Acknowledgments

We thank Quinn Boser, Thomas R. Dawson, Michael R. Dawson, and Albert Vette for experimental design and data processing assistance. This work was supported by the Sensory Motor Adaptive Rehabilitation Technology (SMART) Network at the University of Alberta, the Alberta Machine Intelligence Institute (Amii), and the Canadian Institute for Advanced Research (CIFAR) AI Chairs program, the Killam Trusts, Philanthropic Educational Organization (PEO) International, the Natural Science and Engineering Research Council (NSERC) CGS-D, NSERC DG RGPIN-2019-05961, and NSERC DG RGPIN-2015-03646.

## Declarations

### Competing Interests

The authors have no relevant financial or non-financial interests to disclose. The authors have no competing interests to declare that are relevant to the content of this article. All authors certify that they have no affiliations with or involvement in any organization or entity with any financial interest or non-financial interest in the subject matter or materials discussed in this manuscript. The authors have no financial or proprietary interests in any material discussed in this article.

### Data Availability

All anonymized data supporting the findings of this study will be made available.

## Appendix A Detailed Controller Comparison Results

This appendix outlines results for both controller comparisons, all three tasks, and in all three groups of metrics (Task Performance, Control Characteristics, and User Experience). The two controller comparisons are as follows:

- The recurrent convolutional neural network classification controller (RCNN-Class) versus the linear discriminant analysis baseline classification control (LDA-Baseline)
- The recurrent convolutional neural network regression controller (RCNN-Reg)

versus LDA-Baseline

The controller comparison results are detailed in the three tables below—one table for each group of metrics. Each table displays the median within-participant differences between the controllers in question, calculated as RCNN-Class minus LDA-Baseline or RCNN-Reg minus LDA-Baseline. Interquartile ranges of these differences are presented in parentheses. Green cells indicate metrics in which RCNN-Class or RCNN-Reg performed significantly better than LDA-Baseline (p ¡ 0.05). Grey cells indicate instances where a metric was not relevant. Dark cell borders indicate metrics that displayed evidence of the limb position effect under LDA-Baseline control.

**Table A1.** Task Performance Results.

| Controller Comparison |  | RCNN-Class versus LDA-Baseline |  |  | RCNN-Reg versus LDA-Baseline |  |  |
| --- | --- | --- | --- | --- | --- | --- | --- |
| Task |  | Pasta | RCRT Up | RCRT Down | Pasta | RCRT Up | RCRT Down |
| Success Rate |  | 0.00 (15.00) | 5.00 (10.00) | 5.00 (25.00) | 0.00 (17.50) | 0.00 (17.50) | 0.00 (7.50) |
| Trial Duration (s) |  | -0.93 (10.60) | 1.04 (10.18) | -8.78 (14.40) | -3.03 (11.73) | -2.13 (7.35) | -5.43 (16.64) |
| Phase Duration (s) | Reach | 0.11 (0.76) | -0.03 (1.23) | 0.07 (0.74) | -0.39 (0.75) | -0.63 (0.90) | -0.14 (1.07) |
|  | Grasp | -0.14 (1.01) | -0.12 (1.35) | -0.44 (1.35) | -0.27 (0.98) | -0.37 (1.02) | -0.22 (1.61) |
|  | Transport | -0.03 (0.40) | 0.06 (1.25) | -0.10 (1.40) | -0.34 (0.78) | 0.28 (1.23) | -0.65 (1.43) |
|  | Release | -0.06 (0.41) | 0.20 (1.16) | -0.29 (2.47) | -0.09 (1.03) | 0.05 (1.55) | -0.10 (1.07) |
| Relative Phase Duration (%) | Reach | 0.99 (6.25) | -0.72 (7.30) | 3.79 (12.01) | -1.07 (6.39) | -6.51 (5.46) | 3.11 (9.08) |
|  | Grasp | -2.35 (8.65) | -1.75 (7.68) | -0.37 (12.47) | -2.77 (7.66) | 1.15 (13.75) | 0.99 (8.51) |
|  | Transport | 1.03 (7.17) | -0.47 (8.35) | 2.07 (6.47) | 3.20 (6.39) | 4.15 (5.45) | -1.02 (9.44) |
|  | Release | 0.91 (4.95) | 0.28 (5.32) | -1.73 (12.30) | 0.82 (8.56) | 2.27 (8.17) | 0.16 (8.38) |
| Peak Hand Velocity (mm/s) | Reach-Grasp | 32.71 (163.96) | -21.21 (128.75) | -35.18 (252.92) | 99.01 (335.58) | 111.25 (152.43) | 49.38 (216.65) |
|  | Transport-Release | -20.45 (110.08) | -36.87 (110.66) | -39.07 (129.35) | 117.66 (269.54) | 4.13 (141.52) | 46.48 (102.92) |
| Hand Distance Travelled (mm) | Reach-Grasp | -22.01 (180.61) | 0.26 (84.33) | -16.69 (63.15) | -30.85 (64.24) | -27.17 (50.41) | -6.40 (98.83) |
|  | Transport-Release | -45.52 (70.92) | 10.79 (116.79) | -31.29 (151.25) | -9.99 (41.14) | -1.14 (52.19) | -39.99 (48.94) |
| Hand Trajectory Variability (mm) | Reach-Grasp | -7.85 (43.10) | -3.02 (10.15) | -14.04 (30.23) | -21.77 (30.55) | -7.48 (28.83) | -8.01 (25.14) |
|  | Transport-Release | -17.11 (53.22) | 1.98 (22.24) | -2.54 (21.07) | -6.06 (47.51) | -2.96 (16.29) | -8.54 (15.74) |

**Table A2.**
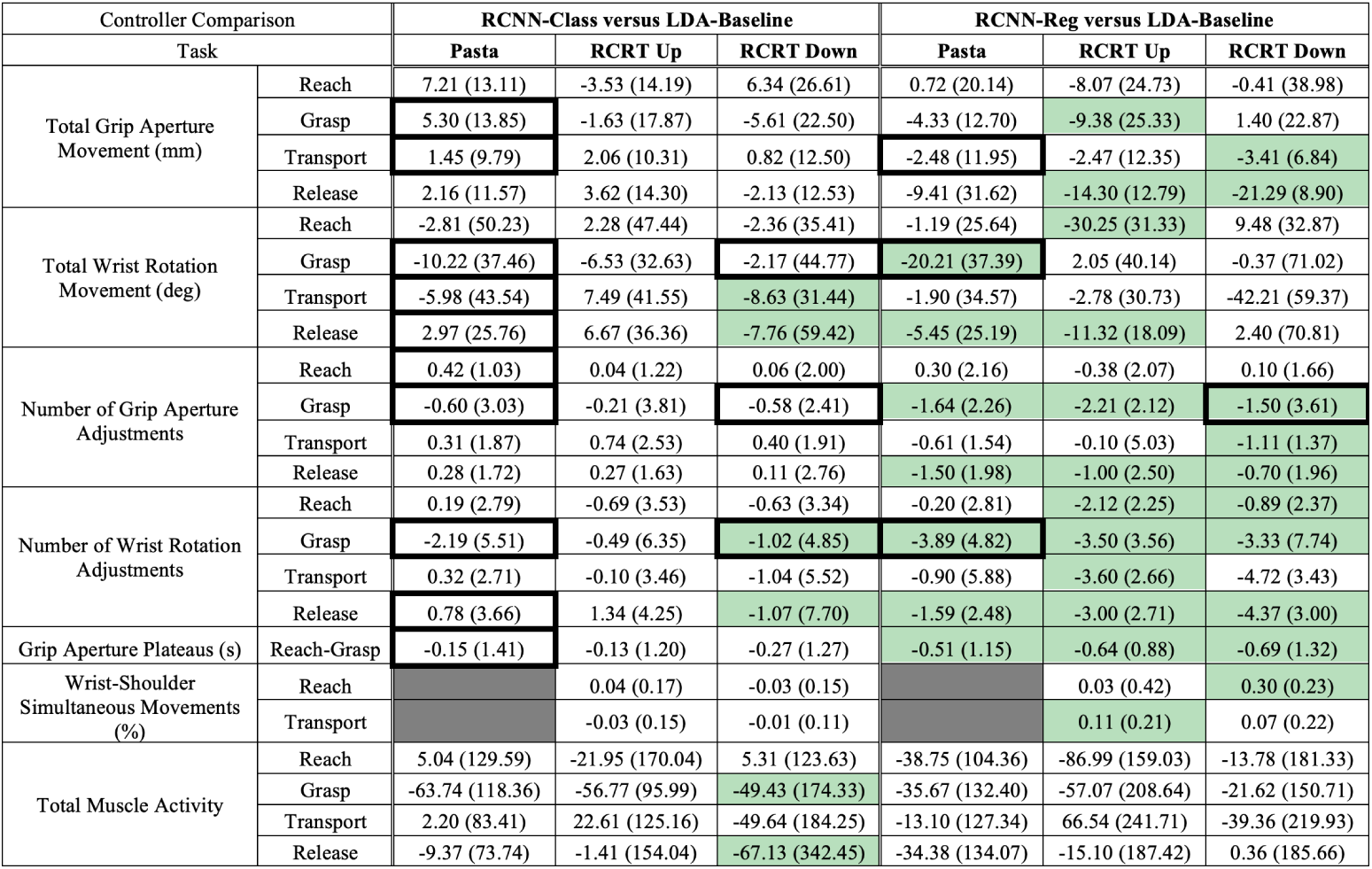
Control Characteristics Results.

| Controller Comparison |  | RCNN-Class versus LDA-Baseline |  |  | RCNN-Reg versus LDA-Baseline |  |  |
| --- | --- | --- | --- | --- | --- | --- | --- |
| Task |  | Pasta | RCRT Up | RCRT Down | Pasta | RCRT Up | RCRT Down |
| Total Grip Aperture Movement (mm) | Reach | 7.21 (13.11) | -3.53 (14.19) | 6.34 (26.61) | 0.72 (20.14) | -8.07 (24.73) | -0.41 (38.98) |
|  | Grasp | 5.30 (13.85) | -1.63 (17.87) | -5.61 (22.50) | -4.33 (12.70) | -9.38 (25.33) | 1.40 (22.87) |
|  | Transport | 1.45 (9.79) | 2.06 (10.31) | 0.82 (12.50) | -2.48 (11.95) | -2.47 (12.35) | -3.41 (6.84) |
|  | Release | 2.16 (11.57) | 3.62 (14.30) | -2.13 (12.53) | -9.41 (31.62) | -14.30 (12.79) | -21.29 (8.90) |
| Total Wrist Rotation Movement (deg) | Reach | -2.81 (50.23) | 2.28 (47.44) | -2.36 (35.41) | -1.19 (25.64) | -30.25 (31.33) | 9.48 (32.87) |
|  | Grasp | -10.22 (37.46) | -6.53 (32.63) | -2.17 (44.77) | -20.21 (37.39) | 2.05 (40.14) | -0.37 (71.02) |
|  | Transport | -5.98 (43.54) | 7.49 (41.55) | -8.63 (31.44) | -1.90 (34.57) | -2.78 (30.73) | -42.21 (59.37) |
|  | Release | 2.97 (25.76) | 6.67 (36.36) | -7.76 (59.42) | -5.45 (25.19) | -11.32 (18.09) | 2.40 (70.81) |
| Number of Grip Aperture Adjustments | Reach | 0.42 (1.03) | 0.04 (1.22) | 0.06 (2.00) | 0.30 (2.16) | -0.38 (2.07) | 0.10 (1.66) |
|  | Grasp | -0.60 (3.03) | -0.21 (3.81) | -0.58 (2.41) | -1.64 (2.26) | -2.21 (2.12) | -1.50 (3.61) |
|  | Transport | 0.31 (1.87) | 0.74 (2.53) | 0.40 (1.91) | -0.61 (1.54) | -0.10 (5.03) | -1.11 (1.37) |
|  | Release | 0.28 (1.72) | 0.27 (1.63) | 0.11 (2.76) | -1.50 (1.98) | -1.00 (2.50) | -0.70 (1.96) |
| Number of Wrist Rotation Adjustments | Reach | 0.19 (2.79) | -0.69 (3.53) | -0.63 (3.34) | -0.20 (2.81) | -2.12 (2.25) | -0.89 (2.37) |
|  | Grasp | -2.19 (5.51) | -0.49 (6.35) | -1.02 (4.85) | -3.89 (4.82) | -3.50 (3.56) | -3.33 (7.74) |
|  | Transport | 0.32 (2.71) | -0.10 (3.46) | -1.04 (5.52) | -0.90 (5.88) | -3.60 (2.66) | -4.72 (3.43) |
|  | Release | 0.78 (3.66) | 1.34 (4.25) | -1.07 (7.70) | -1.59 (2.48) | -3.00 (2.71) | -4.37 (3.00) |
| Grip Aperture Plateaus (s) | Reach-Grasp | -0.15 (1.41) | -0.13 (1.20) | -0.27 (1.27) | -0.51 (1.15) | -0.64 (0.88) | -0.69 (1.32) |
| Wrist-Shoulder Simultaneous Movements (%) | Reach |  | 0.04 (0.17) | -0.03 (0.15) |  | 0.03 (0.42) | 0.30 (0.23) |
|  | Transport |  | -0.03 (0.15) | -0.01 (0.11) |  | 0.11 (0.21) | 0.07 (0.22) |
| Total Muscle Activity | Reach | 5.04 (129.59) | -21.95 (170.04) | 5.31 (123.63) | -38.75 (104.36) | -86.99 (159.03) | -13.78 (181.33) |
|  | Grasp | -63.74 (118.36) | -56.77 (95.99) | -49.43 (174.33) | -35.67 (132.40) | -57.07 (208.64) | -21.62 (150.71) |
|  | Transport | 2.20 (83.41) | 22.61 (125.16) | -49.64 (184.25) | -13.10 (127.34) | 66.54 (241.71) | -39.36 (219.93) |
|  | Release | -9.37 (73.74) | -1.41 (154.04) | -67.13 (342.45) | -34.38 (134.07) | -15.10 (187.42) | 0.36 (185.66) |

**Table A3.** User Experience Results.

| Controller Comparison |  | RCNN-Class versus LDA-Baseline |  |  | RCNN-Reg versus LDA-Baseline |  |  |
| --- | --- | --- | --- | --- | --- | --- | --- |
| Task |  | Pasta | RCRT Up | RCRT Down | Pasta | RCRT Up | RCRT Down |
| NASA-TLX | Mental Demand | -10.00 (22.50) | -10.00 (22.50) | -10.00 (22.50) | 20.00 (22.50) | 20.00 (22.50) | 20.00 (22.50) |
|  | Physical Demand | -10.00 (27.50) | -10.00 (27.50) | -10.00 (27.50) | 5.00 (23.75) | 5.00 (23.75) | 5.00 (23.75) |
|  | Temporal Demand | 5.00 (22.50) | 5.00 (22.50) | 5.00 (22.50) | -10.00 (41.25) | -10.00 (41.25) | -10.00 (41.25) |
|  | Performance | -5.00 (12.50) | -5.00 (12.50) | -5.00 (12.50) | -5.00 (13.75) | -5.00 (13.75) | -5.00 (13.75) |
|  | Effort | -2.50 (27.50) | -2.50 (27.50) | -2.50 (27.50) | 0.00 (26.25) | 0.00 (26.25) | 0.00 (26.25) |
|  | Frustration | -5.00 (40.00) | -5.00 (40.00) | -5.00 (40.00) | -5.00 (13.75) | -5.00 (13.75) | -5.00 (13.75) |
| Usability | Intuitiveness | 0.40 (0.95) | 0.40 (0.95) | 0.40 (0.95) | 0.00 (0.88) | 0.00 (0.88) | 0.00 (0.88) |
|  | Effectiveness - RCRT | -0.25 (0.85) | -0.25 (0.85) | -0.25 (0.85) | 0.62 (1.15) | 0.62 (1.15) | 0.62 (1.15) |
|  | Effectiveness - Pasta | 0.40 (1.25) | 0.40 (1.25) | 0.40 (1.25) | 0.62 (1.17) | 0.62 (1.17) | 0.62 (1.17) |
|  | Reliability | 0.00 (1.40) | 0.00 (1.40) | 0.00 (1.40) | 0.38 (2.02) | 0.38 (2.02) | 0.38 (2.02) |

## Notes

### Competing Interest Statement

The authors have declared no competing interest.

